# Effects of sex and gestational exercise on pain perception, BDNF and irisin levels in an animal model of ADHD

**DOI:** 10.1101/2023.08.04.551984

**Authors:** Andréa Tosta, Ariene S. Fonseca, Débora Messender, Sérgio T. Ferreira, Mychael V. Lourenco, Pablo Pandolfo

## Abstract

Abnormal cognitive and sensorial properties have been reported in patients with psychiatric and neurodevelopmental conditions, such as attention deficit hyperactivity disorder (ADHD). ADHD patients exhibit impaired dopaminergic signaling and plasticity in brain areas related to cognitive and sensory processing. The spontaneous hypertensive rat (SHR), in comparison to the Wistar Kyoto rat (WKY), is the most used genetic animal model to study ADHD. Brain neurotrophic factor (BDNF), critical for midbrain and hippocampal dopaminergic neuron survival and differentiation, is reduced in both ADHD subjects and SHR. Physical exercise (e.g. swimming) promotes neuroplasticity and improves cognition by increasing BDNF and irisin. Here we investigate the effects of gestational swimming on sensorial and behavioral phenotypes, striatal dopaminergic parameters, and hippocampal FNDC5/irisin and BDNF levels observed in WKY and SHR. Gestational swimming improved nociceptive reflex impairment in SHR rats and increased hippocampal BDNF levels in a sex-dependent manner in adolescent offspring. Sex differences were observed in hippocampal FNDC5/irisin levels, with females presenting lower levels than males. Our results contribute to the notion that swimming during pregnancy is a promising alternative to improve ADHD phenotypes in the offspring.

**Graphical abstract:** 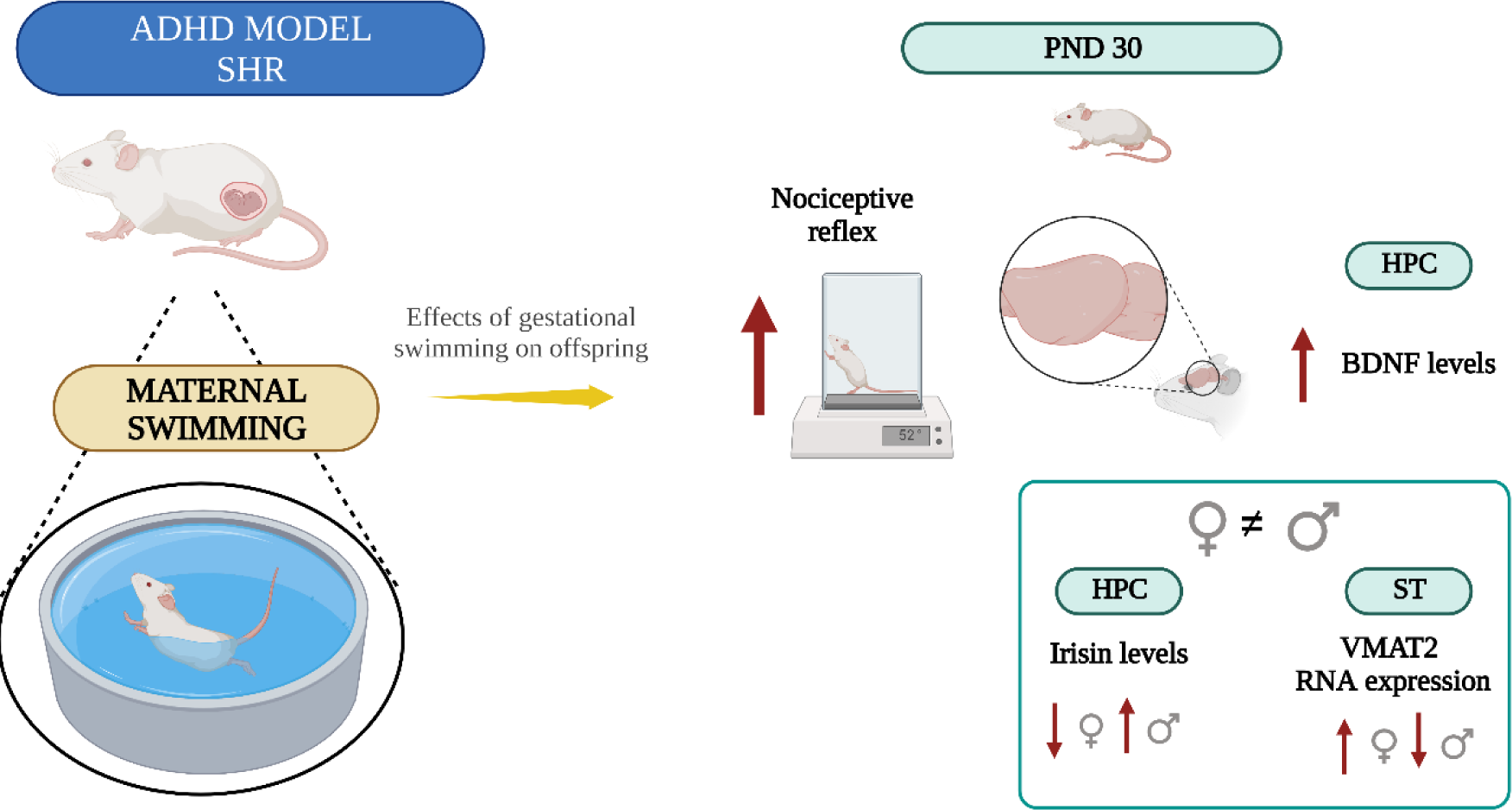

## 1. Introduction

Attention deficit hyperactivity disorder (ADHD) is a neurodevelopmental disorder with a higher prevalence among young males compared to females (Skogli et al., 2013). It is characterized by cognitive deficits, dysregulation of sensory processing, and delayed plasticity in brain areas such as the striatum and hippocampus (Biederman and Faraone, 2005; Faraone et al., 2015; Polanczyk and Rohde, 2007). Abnormal pain experiences have been reported in patients with ADHD (Johnston and Huckins, 2022; Lautenbacher and Krieg, 1994). Pain perception involves both the attentional state and emotional profile processed in brain limbic areas (Becker et al., 2023; Bouchatta et al., 2022). Functional changes in frontostriatal monoaminergic circuits are found in ADHD and chronic pain (Faraone, 2018; Zhang et al., 2017). Modulation of monoaminergic neurotransmission is a pharmacological target for these conditions managed with methylphenidate or antidepressants (Treister et al., 2015).

The strain pair spontaneous hypertensive rat (SHR) and Wistar Kyoto rat (WKY) comprise the most used genetic animal model to study ADHD (Rahi and Kumar, 2021). In addition to showing neurobiological and behavioral germane to ADHD (Rahi and Kumar, 2021), SHR present abnormal nociceptive reactivity (Pamplona et al., 2007; Vendruscolo et al., 2004).

Evidence suggests that dopamine is involved not only in core ADHD symptoms but also in sensorial deficits, such as pain perception (Johnston and Huckins, 2022; Wolff et al., 2016). The mesolimbic dopaminergic pathway plays a crucial role in locomotor activity, attentional behavior, and pain sensation (Mehta et al., 2019; Miller et al., 2012; Wood, 2006). The hippocampus is further targeted by dopaminergic neurons of the mesolimbic pathway (Gasbarri et al., 1994), also altered in ADHD individuals.

Brain-derived neurotrophic factor (BDNF) is critical for midbrain dopaminergic neuron survival and differentiation (Knosel et al., 1991; Liu et al., 2015). Midbrain BDNF signaling also impacts the dopaminergic neurotransmission underlying ADHD and neuropathic pain (Liu et al., 2015; Zhang et al., 2017; Marques et al., 2023). Physical exercise promotes hippocampal expression of BDNF (Gómez-Pinilla et al., 2002; Lourenco et al., 2019a) and irisin, an exercise-induced hormone that mediates synaptic plasticity and memory (Abdulghani et al., 2023; Isaac et al., 2021; Lourenco et al., 2019a; Wrann et al., 2013). Furthermore, exercise increased striatal dopamine release through BDNF ((Bastioli et al., 2022)) and stimulates monoaminergic synapses (Liu et al., 2015).

Exercising during pregnancy promotes benefits for both mother and offspring (Clapp, 1996). Studies in animal models have shown that maternal swimming benefits the offspring, which includes increased hippocampal plasticity, prevents mitochondrial dysfunction, improves learning and memory, and promotes short- and long-term neuroprotection (Klein et al., 2020; Lee et al., 2006; Marcelino et al., 2016; Sanches et al., 2020; Tosta et al., 2023). Here we investigate the effects of gestational swimming on pain perception, cognition, motor coordination, dopaminergic parameters, and BDNF and irisin levels in an ADHD animal model.

## 2. Materials and methods

### 2.1. Experimental subjects

Male and female isogenic rats of approximately 90 days (WKY and SHR strains) were used for mating. They were obtained from the Multidisciplinary Center Biological Research in the Area of Science in Laboratory Animals (CEMIB) of the State University of Campinas (UNICAMP) (Campinas, SP, Brazil) and kept in the sectoral vivarium of the Department of Neurobiology, Institute of Biology of the University Federal Fluminense (Niterói, RJ, Brazil). They were maintained on standard food and water *ad libitum* regimen, and 12-hour light/dark cycles in room with controlled temperature (24°C ± 2°C). WKY and SHR were placed to mate in Plexiglass cages (42x34x18 cm), distributed as follows: two females and one male per cage during mating. All animal experiments were performed following the principles of the “3Rs” and the National Institutes of Health Guide for the care and use of laboratory animals.

### 2.2. Experimental design

After pregnancy confirmation, rats were immediately subjected to the swimming protocol, from gestational day (GD) 0 to GD20. Swimming was performed five times a week for 20 minutes. On the twentieth day of gestation, females were placed in isolated cages, remaining until parturition. On the post-natal day (PND) 7 and PND14, the offspring was weighed. Motor reflexes were evaluated in the PND14. On PND 21, the animals were weighed, weaned, and placed in cages according to sex, strain, and treatment. Behavioral tasks performed: Y-maze (YM) (PND30), rotarod habituation (PND31), rotarod test (PND32), and nociceptive reflex (PND33). Twenty-four hours after the last behavioral tasks (PND34), the striatum and hippocampus were collected for biochemistry analyses (Fig. 1).

**Figure 1.**
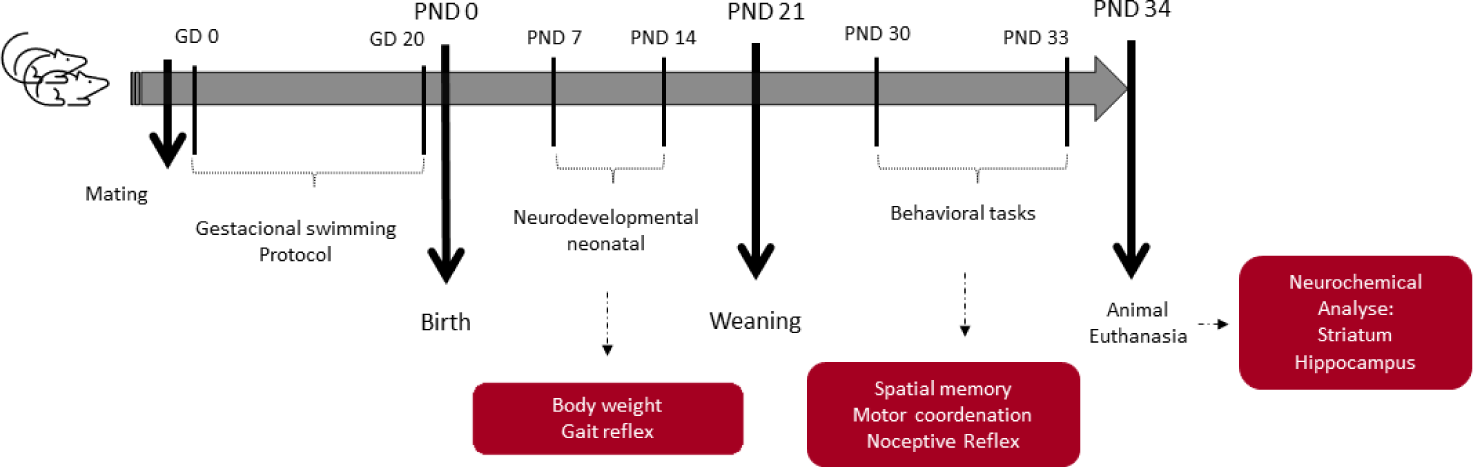
Schematic overview of the experimental design. 90-day-old male and female rats of strains WKY and SHR were mated. After confirmation of pregnancy, the rats were immediately subjected to the swimming protocol, on the gestational day (GD) 0, lasting until GD20, swimming was performed five times *per* week for 20 continuous minutes. On the twentieth day of gestation, females were housed in isolated boxes and remained there until parturition. The day of birth is considered postnatal day (PND) 0. At PND 7 and PND 14, animals were weighed and SHIRPA tests were performed to analyze the development and sensorimotor reflexes of the offspring. In PND 21 the animals were weaned and placed in boxes according to sex, strain, and treatment. From PND 30, behavioral tests were performed during adolescence: spatial memory in the Y-maze (YM), motor coordination in the rotarod (RR) and nociceptive reflex in the hot plate (HP). Twenty-four hours after the final behavioral tasks, PND 34, animals were anesthetized with isoflurane, decapitated, and striatum and hippocampus were collected for biochemical analyzes.

### 2.3. Swimming exercise protocol

The swimming apparatus consisted of a circular plastic tank (50 x 30 cm) filled with water at 32 ± 1°C (adapted from Lee et al. 2006). The swimming training protocol consisted of daily free-swimming sessions of 20 min/5 times a week from the first to the twentieth day of pregnancy. Females were divided into 4 groups: 2 control groups (sedentary groups: WKY SED and SHR SED) and 2 exercised groups (swimming groups: WKY SWIM and SHR SWIM). The control groups, WKY SED and SHR SED, were only exposed to the training environment: a tank filled with water at a shallow level approximately 1 cm deep for 5 min. After the last training session, each pregnant rat was housed alone in a clean standard cage until parturition.

### 2.4. Body weight and gait reflex

The gait reflex test was performed in a horizontal apparatus in which a circle approximately 13 cm in diameter is drawn. The latency for the animal to move away from the center of the circle until the entire body is outside the circle was measured, the maximum time being 60 seconds (Cárdenas et al., 2015; Sanches et al., 2017a).

### 2.5. Behavioral tasks in adolescence

Animals were acclimated to the experimental room for one hour prior to both neurodevelopmental and behavioral tests. Behaviors tasks were performed in an enclosed, quiet room with low-intensity indirect lighting. Equipment and objects were cleaned with 10% alcohol between trials to avoid odor traces. Animal behavior was recorded with a camera mounted above the apparatus and quantified using AnyMaze® software.

#### Y-maze (EPM)

Rat spatial memory was assessed using Y-maze (Dellu et al., 1992; Pandolfo et al., 2013). The test was performed in an apparatus with three equal arms (50 x 10 x 2043 cm, each arm) arranged at an angle of 120°. The room in which it was performed had visual cues on the walls that facilitated spatial location. The test was performed in two trials separated by an one-hour interval. In the first trial, access to one of the arms was blocked and the animal had free access to the other two arms for 5 minutes. The arm that was blocked was designated as the new arm, and in the second trial, one hour later, the animal was allowed to explore the three arms for 5 minutes. Results were represented by the percentage of entries and time spent in the new arm.

#### Rotarod

Motor coordination and balance were assessed using the rotarod accelerator (Qian et al., 2010). The protocol was divided in two days, with the first day considered the adaptation period to the device. The animal was placed in the apparatus at low and constant speed (4 rpm), 3 trials of 1 minute were performed with an interval of 5 minutes between trials. The test was performed 24 hours after training, where the animal’s ability to maintain balance while walking on the apparatus was measured. Three 5-minute trials were performed with a 5-minute intertrial interval. The rotation speed varied from 4 to 40 rpm. Therefore, the fall latency of each subject was analyzed through the mean of the 3 trials.

#### Nociceptive reflex

One hour after the motor learning test, the animals were subjected to the nociceptive reflex test. A hot plate with a temperature of 52°C ± 1°C was used to measure the nociceptive sensitivity of the animals. Rats were placed under the hot plate, and the first sign of discomfort (licking, jumping, vocalizing, or lifting the paws from the hot plate), the rat was immediately removed from the hot plate, and the latency of response was recorded (Ankier and Ankier, 1974).

### 2.6. Western blotting

Twenty-four hours after the last behavioral test, rats were euthanized, and the striatum and hippocampus were collected. The striatal and hippocampal samples were stored at -80°C until homogenization in RIPA buffer (25 mM Tris-HCl pH 7.6, 150 mM NaCl, 1% NP-40, 1% sodium deoxycholate, 0.1% SDS; Thermo Fisher) containing cocktails of protease and phosphatase inhibitors. Samples were resolved in pre-cast 4- 20% polyacrylamide gels (BioRad®) with Tris/glycine/SDS buffer and resolved at 100 V for 90 min at room temperature. The gel (30 µg total protein/lane) was electroblotted onto Hybond ECL nitrocellulose using 25 mM Tris, 192 mM glycine, 20% methanol (v/v), pH 8.3, at 350 mA for 60 min at 4°C. Membranes were blocked with a 3% BSA in TBS-T for 1 hour at room temperature. The membranes were incubated under agitation with primary antibodies (anti-DAT, anti-TH, anti-D_2_R (1:1000, diluted in TBS-T and 3% BSA), anti-β-actin monoclonal antibodies (1:2000, diluted in TBS-T and 3% BSA) (Supplementary Table 1) overnight at 4°C. After incubation with secondary anti-rat and anti-mouse IgGs (1:10,000, diluted in TBS-T and 3% BSA) (Supplementary Table 1) for 120 min, membranes were washed and developed in Odyssey. The images obtained were analyzed using the Image J program to obtain the average optical density values of the bands.

### 2.7. ELISA

Hippocampal tissue was collected and centrifuged at 10,000 × g for 10 min at 4°C and the supernatant was supplemented with protease and phosphatase inhibitor cocktails. BDNF assays were performed according to the kit manufacturer’s instructions (Abcam, ab212166), after sample dilution optimization.

### 2.8. Quantitative RT-PCR

Total RNA was extracted from striatum samples of adolescent WKY and SHR animals using Trizol® (Invitrogen), following the manufacturer’s instructions. Isolated total RNA was eluted in 20 µL of nuclease-free water. RNA concentration and purity were determined with Nanodrop (Biochrom) to calculate the optical density ratio at the wavelength of 260/280 nm and 260/230 nm. For qRT-PCR, 1 µg of total RNA was used for cDNA synthesis using the High-Capacity cDNA Reverse Transcription Kit (Applied Biosystems, CA). Quantitative analysis of target gene expression was performed on a 7500 Applied Biosystems real-time PCR system (Foster City, CA) with the Power SYBR Green master mix. β-actin (actb) was used as an endogenous reference gene for data normalization. qRT-PCR was performed in reaction volumes of 12 µL according to the manufacturer’s protocols. The primer sequences used for qPCR are described in Supplementary Table 2. Cycle threshold (Ct) values were used to calculate fold changes in gene expression using the 2^-ΔΔCt^ method (Livak and Schmittgen, 2001).

### 2.9. Statistical analyses

Data were expressed as mean ± standard error of the mean (SE). Three-way analysis of variance ANOVA (treatment x strain x sex) was performed to analyze offspring weight gain at PND 7 and PND 14, neonatal neurodevelopmental reflexes, and all behavioral tests and biochemical analyzes. When results were significant at ANOVA, multiple comparisons were performed using the Tukey post hoc multiple frequency test (Tukey test for multiple comparisons). The significance level used in all experiments was p ≤ 0.05. Statistical analyzes were performed using JASP software (Jeffreys’s Amazing Statistics Program, Amsterdam) and graphs were generated using Prism 8.0 software (Graphpad®).

## 3. Results

### 3.1. Sex differences in body weight of strains and effects of maternal swimming on weight and gait reflex

Weighing was performed on PND7 and PND14 to evaluate body mass gain in the offspring. On PND7, the control strain WKY had higher body mass than the strain SHR (three-way ANOVA, strain factor: F (1, 16) = 126.4, p ≤ 0.05). The higher weight of males was also observed compared to females, regardless of strain (three-way ANOVA, sex factor: F (1, 16) = 10.8, p ≤ 0.05). Gestational swimming did not alter weight on PND7 (Fig. 2A). On PND14 (Fig. 2B), the difference in weight between WKY and SHR strains persisted (three-way ANOVA, strain factor: F (1, 16) = 211.8, p ≤ 0.05), as did the difference between sexes (ANOVA three-way, sex factor: F (1, 16) = 12.7, p ≤ 0.05). On the other hand, selectively in the WKY group, the animals whose mothers swam had more weight gain compared to those from sedentary mothers (three-way ANOVA, interaction between strain vs. treatment: F (1, 16) = 29.7, p ≤ 0.05).

**Figure 2.**
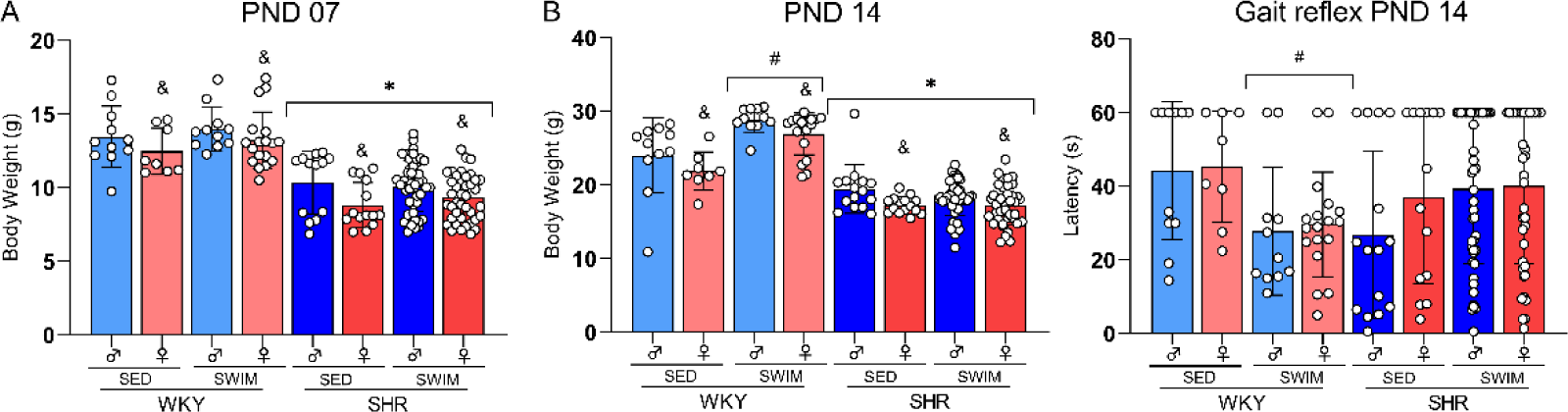
Effect of gestational swimming on neonatal neurodevelopment of WKY rat offspring. Comparison between offspring weight gain in A) PND 7 (n= 8-46 animals *per* group) and in B) PND 14 (n= 8-46 animals *per* group), and C) gait reflex (n=8-46 animals *per* group). Data are presented as mean ± SEM ^&^p < 0.05 male vs. female, regardless of treatment or strain; ^#^p <0.05 SED vs. SWIM treatments in the same strain, regardless of sex; *p <0.05 WKY vs. SHR strains, regardless of treatment and sex (three-way ANOVA; *post hoc* Tukey).

On PND 14, the motor reflex test was performed after weighing the animals (Fig. 2C). Analysis of the results showed an interaction between strain vs. treatment (three-way ANOVA, F (1, 16) = 10.5, p ≤ 0.05). Regardless of sex, WKY SWIM took less time on average to fully exit the diameter than the SED group. This indicates an effect of gestational swimming on the motor reflex of the WKY offspring.

### 3.2. Swimming during pregnancy reverses the impairment of pain perception in adolescent SHR

On PND 30, behavioral tasks assessing spatial memory, motor coordination and nociceptive reflex were performed (Fig. 3). Results showed a strain effect on spatial memory in the number of entries (three-way ANOVA: F (1, 93) = 36,5, p < 0.001) (Fig. 3A) and exploration time (three-way ANOVA: F (1, 92) = 5,4, p ≤ 0.05) in the new arm (Fig. 3B). SHR, regardless of treatment and sex, spent less time and made less entries in the novel arm than the WKY control group. No difference was observed for the strains, sexes, or treatment in the motor coordination test (Fig. 3C).

**Figure 3.**
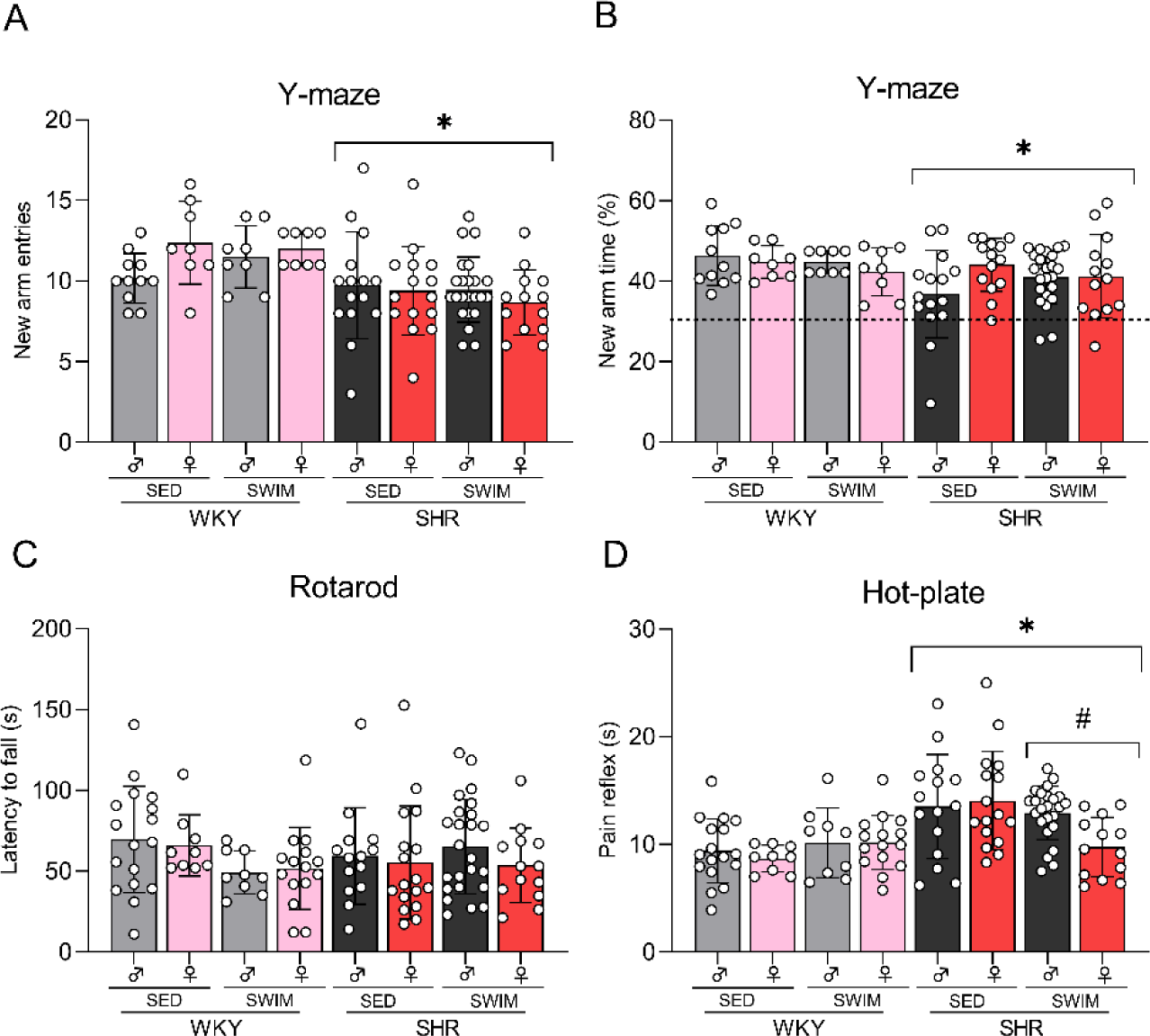
Behavioral tests in adolescence. Bars show mean ± SEM in (A) number of entries into new arm (%) n=8-23 animals *per* group, (B) total time in new arm (%), n=8-23 animals *per* group, (C) average rotarod fall latency n=8-23 animals *per* group and (D) latency of pain perception on hot plate n= 8-23 animals *per* group *p<0.05 compared with the WKY strain, regardless of treatment or sex, ^#^p<0.05 compared with the same-sex, same-strain SED group (three-way ANOVA; post hoc Tukey).

Nociceptive sensitivity was measured 1 hour after the rotarod test, on the third day of the behavioral battery. The nociceptive stimulus latency of WKY and SHR strains exposed or not to maternal swimming is shown in Figure 3D. The three-way ANOVA showed a significant difference for the strain factor (F (1,11) = 21.6, p ≤ 0.05). The post-hoc test showed that SHR animals took longer to perceive pain than control animals regardless of treatment and sex. This result is consistent with the literature showing a delay in nociceptive sensitivity in rats SHR, a phenotype also observed in patients with ADHD. In addition, results showed the effect of swimming during pregnancy on the interaction between strain vs. treatment (F (1,11) = 7.8, p ≤ 0.05). Therefore, adolescent SHR whose mothers swam had a faster nociceptive reflex than sedentary mothers.

### 3.3. Analysis of the protein content of the dopaminergic system in the striatum

Protein content of DAT (Fig. 4A and 4B), TH (Fig. 4C and 4D), and D_2_R (Fig. 4E and 4F) was measured in the striatum of WKY and SHR. Three-way ANOVA showed no significant differences in the protein levels examined.

**Figure 4.**
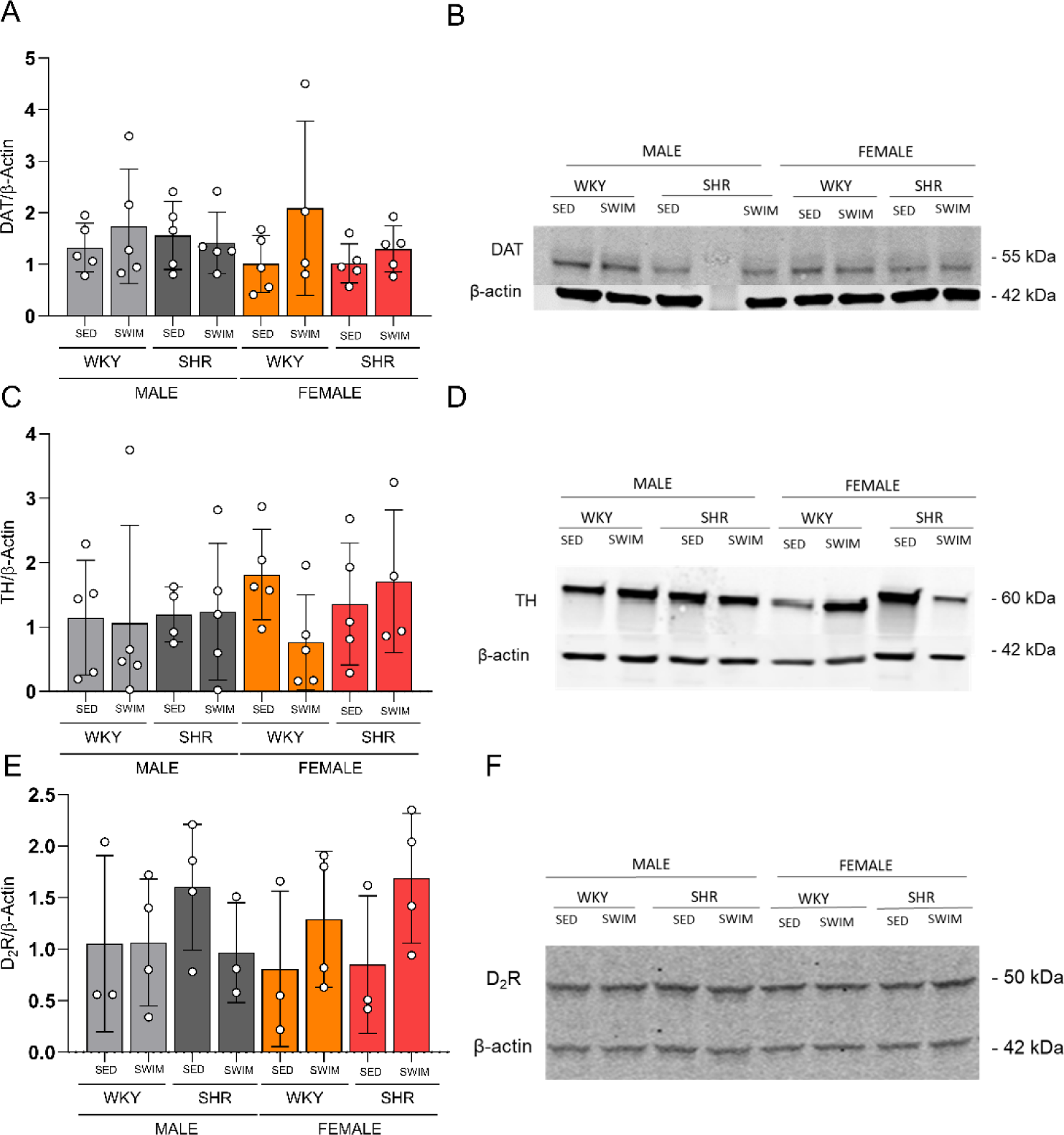
Analysis of expression density of dopaminergic parameters in the striatum. Bars represent the means ± SEM (n=4 animals per group) of density of (A) dopamine transporters (DAT), (C) tyrosine hydroxylase enzyme (TH) and (E) dopaminergic D2 receptors (D2DR). Representative blots (B, D, F) in samples of the striatum of WKY and SHR rats exposed or not to maternal swimming (SWIM and SED).

### 3.4. Strain and sex-dependent effects on striatal D2R and vesicular monoamine transporter 2 (VMAT2) expression levels in adolescent rats SHR

RNA content in the striatum was analyzed for TH (Fig. 5A), DAT (Figure 5B), D_2_R (Figure 5C), and VMAT2 (Fig. 5D). Three-way analysis ANOVA showed a strain effect [F (1,30) = 11.56, p ≤ 0.001] on D_2_R expression. SHR showed higher D_2_R expression compared to WKY animals. Analysis of VMAT2 expression level showed an interaction between strain vs. sex [F (1,30) = 4.534, p = 0.042]. SHR females showed higher VMAT2 expression than males, regardless of treatment.

**Figure 5.**
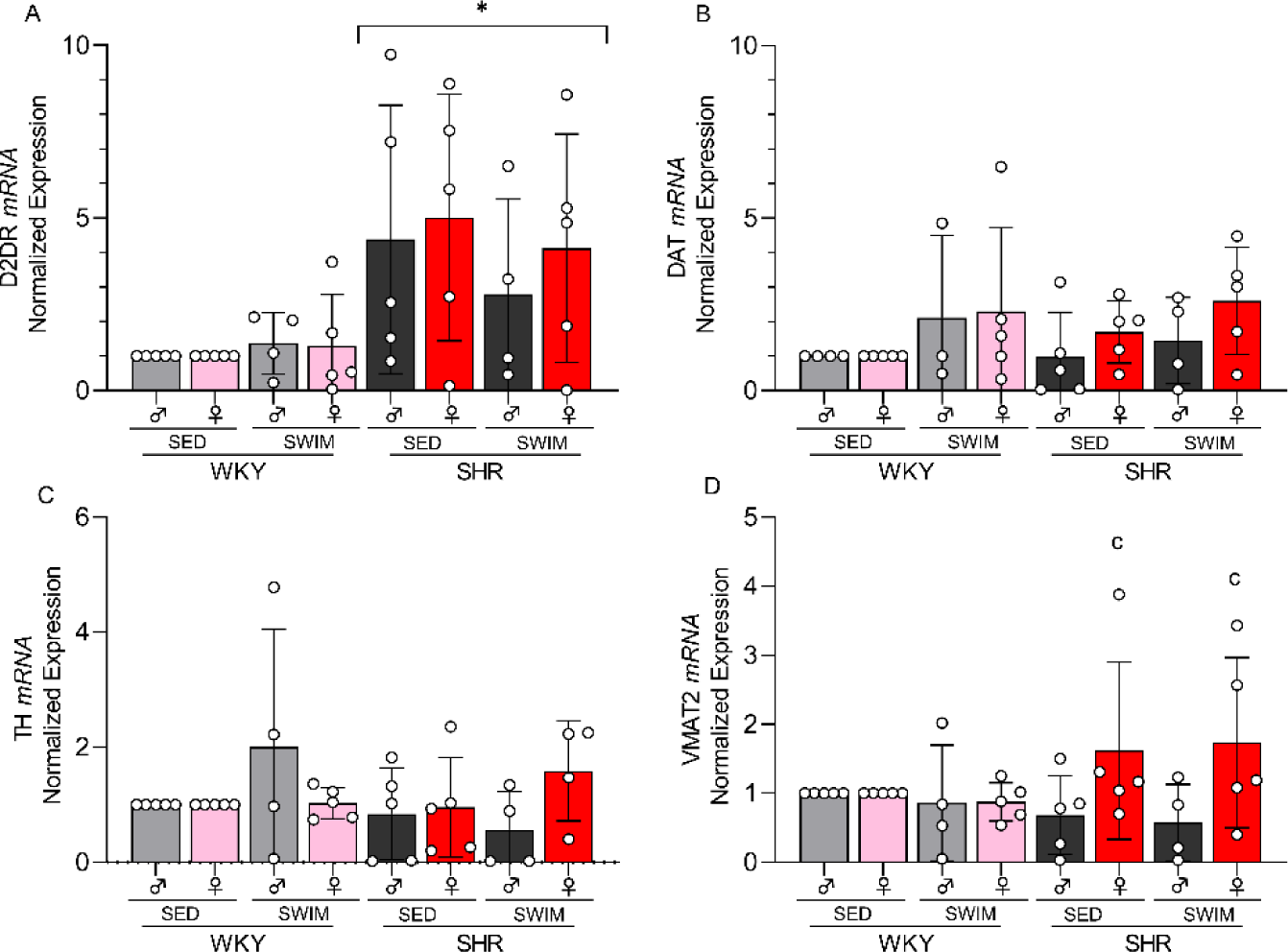
Analysis of expression of dopaminergic parameters in the striatum. Bars represent the means ± SEM (n=3-5 animals per group) of expression of (A) dopaminergic D2 receptors (D2DR), (B) dopamine transporters (DAT), (C) tyrosine hydroxylase enzyme (TH) and (D) vesicular monoamine transporter 2 (VMAT2) in samples of the prefrontal cortex of WKY and SHR rats exposed or not to maternal swimming (SWIM and SED). *p<0.05 WKY vs. SHR strains, regardless of treatment and sex; ^#^p<0.05 SED vs. SWIM treatments in the same strain, regardless of sex (three-way ANOVA; *post hoc* Tukey).

**Figure 6.**
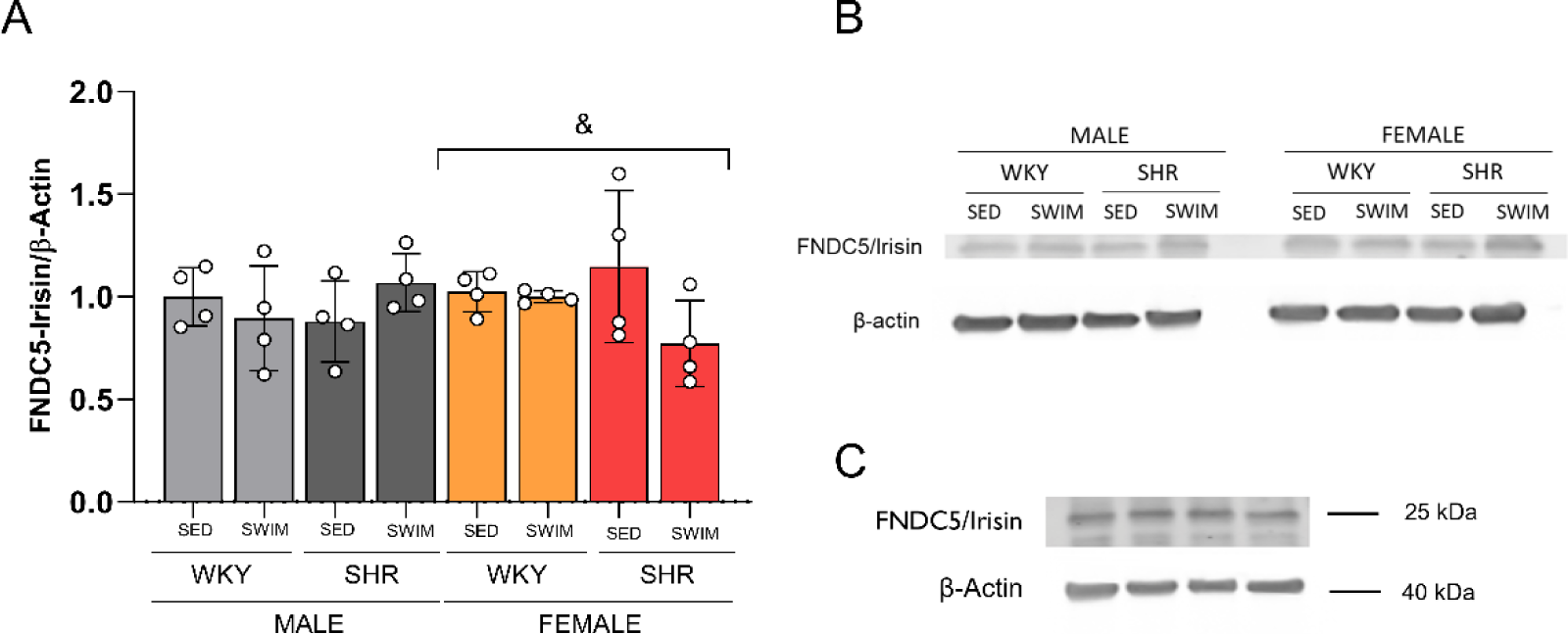
Effect of sex on hippocampal irisin levels. Bars show mean ± SEM expression of FNDC5/Irisin n= 4 animals *per* group. ^&^p<0.05 compared with MALE and FEMALE, same-strain and regardless of treatment (three-way ANOVA; post hoc Tukey).

### 3.5. Sex-related difference in neuroprotective parameters in the hippocampus

As shown in Figure 7, three-way ANOVA, revealed that irisin levels had a sex-dependent effect [F (1,24) = 5.722, p = 0.002]. Female rats had lower irisin levels than males regardless of strain and treatment. No effect of maternal swimming on irisin levels was observed. However, an interaction between strain, treatment, and sex was observed when BDNF levels were assessed in the hippocampus (Figure 8) [F (1,55) = 5.076, p = 0,02]. The SHR strain had lower BDNF levels than the WKY control group, and, interestingly, maternal swimming reversed this effect in the SHR males compared to females in the SWIM group and the SED group, regardless of sex, thereby suggesting that gestational exercise may rescue hippocampal BDNF levels in the SHR male offspring.

**Figure 7.**
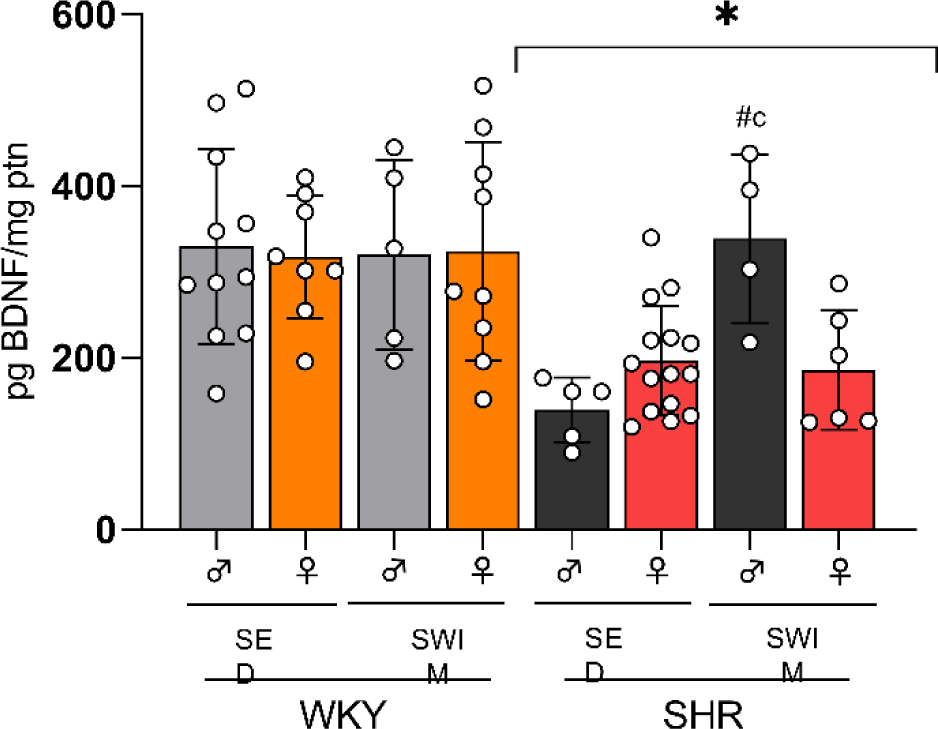
Gestational swimming increases hippocampal BDNF levels in males SHR. Bars show mean ± SEM expression of BDNF. n= 4-15 animals *per* group. *p<0.05 compared with WKY strain, regardless of treatment or sex, ^#^p<0.05 compared with same-sex, same-strain SED group, ^&^p<0.05 compared with male and female, same-strain and regardless of treatment (three-way ANOVA; post hoc Tukey) (three-way ANOVA; post hoc Tukey).

## 4. Discussion

In this study, we investigated the effects of swimming during pregnancy on behavioral, dopaminergic, and neuroprotective parameters in the offspring of an animal model of ADHD. SHR rats exhibited lower body mass in the first two postnatal weeks, memory impairment, delayed pain response, lower D_2_R levels in the striatum, and lower levels of BDNF in the hippocampus compared to WKY rats. Gestational swimming improved the nociceptive reflex impairment and increased BDNF levels in a sex-dependent manner in SHR rats. Sex differences were also observed in the lower levels of vesicular monoamine transporter 2 (VMAT2) in the striatum and higher levels of FNDC5/irisin in the hippocampus of males compared to females SHR.

The notion that exercise has beneficial effects on human health as a non-pharmacological therapeutic approach is widely recognized. In ADHD children and animal models, exercise improves behavioral symptoms and promotes positive changes in the dopaminergic system (Archer and Kostrzewa, 2012; Jeong et al., 2014; Ko et al., 2013; Pontifex et al., 2013; Robinson et al., 2011). Regular physical exercise increases cerebral blood flow, improves cognitive functions including memory and attention, and neuronal plasticity and neuroprotection (de Freitas et al., 2020; Isaac et al., 2021; Lourenco et al., 2022, 2019; Praag, 2008; Praag et al., 2005). In addition, studies in animal models suggest that physical exercise may have intergenerational effects on brain health and development (Valkenborghs et al., 2022). Therefore, exercising during pregnancy has been increasingly recommended by physicians (Medical Association, 2017). We thus examined whether gestational swimming in SHR rats, an animal model of ADHD, promoted neuroprotection from neonatal to adolescence.

SHR have been reported to present lower body mass than their WKY control (Ferguson et al., 2003). Weighings were performed on PND7 and PND14, and on both timepoints, a difference in weight between the strains was observed. We found that SHR neonates have a slower motor reflex as compared to WKY neonates. This may be related to the maturational delay presented in both individuals and ADHD animal models (Ferguson et al., 2003; Sanches et al., 2017). Sex differences were also assessed. As predicted, males of both strains had higher body mass than females. In the SHR strain, differences are also persistent, with females tending to be smaller and more active than males (Berger and Sagvolden, 1998).

We also evaluated the effects of maternal swimming during adolescence. SHR are reported to have spatial memory deficits (Nunes et al., 2018a; Pandolfo et al., 2013; Sontag et al., 2013) and some authors attribute this low performance to the hyperactivity presented by this strain, generating a confounding factor (Nunes et al., 2018b; Sontag et al., 2013). The results corroborate the difference between the SHR and WKY strains in the Y-maze test (Pandolfo et al., 2013). Although the two strains had rates above 33%, which are supportive of task learning, SHR had impaired short-term spatial memory compared to WKY. Gestational swimming did not influence memory impairment in SHR.

Problems in motor performance and coordination also occur in patients with the disorder and are related to excessive movements, lack of balance, difficulty learning and performing certain motor skills, such as tying shoes and playing sports, and impaired fine motor skills (Qian et al., 2010). Quian et al. (2010) performed the rotarod test in Wistar and SHR rats and did not observe a significant difference between the strains. Likewise, the presented result showed no difference between the strains, confirming that SHR rats do not present deficits in motor coordination. Thus, it is believed that SHR rats have deficits in fine motor skills (Qian et al., 2010), which could not be evaluated in the rotarod test.

ADHD is a neurodevelopmental disorder with associated behavioral, emotional, and cognitive deficits. Both ADHD and pain are complex and have a multifactorial origin. Evidence shows that modulation of pain promotes a less intense perception when individuals are distracted by a noxious stimulus (Bushnell et al., 1999; Levine et al., 1982; Miron et al., 1989; Rode et al., 2001). Studies show that SHR have nociceptive impairment in some nociceptive tests (Vendruscolo et al., 2004), including the hot plate test when compared to rats of other strains, such as WKY (Pamplona et al., 2007). However, reports show that they have nociceptive fibers with their normal properties (Bulka and Wiesenfeld-Hallin, 2003). Therefore, evidence indicates that the hypoalgesia in SHR may involve an attentional component (Pamplona et al., 2007). Interestingly, gestational swimming reversed nociceptive reflex damage with a decrease in pain perception time in SHR rats. However, future studies are warranted to better understand pain modulation in SHR.

The neurochemical hypothesis of ADHD is associated with dopaminergic dysfunction in the cortico-striatal regions (Faraone et al., 2015). The striatum is a major source of dopamine in the brain (Hagelberg et al., 2004; Mehta et al., 2019), suggesting an important role for dopamine in attentional processes (van der Kooij and Glennon, 2007). Dopamine is thought to control movement, cognition, perception of pain, and reward (Ohno, 2003). The dopamine D_2_ receptor plays an important role in dopamine regulation and signaling (Ohno, 2003). The expression of the dopamine D_2_ receptor in the striatum was increased in SHR, a result well described in the literature (Cho et al., 2014). The role of VMAT2 is to protect monoamines, including dopamine, from mitochondrial oxidation of MAO in the cytoplasm (Simchon et al., 2010). A study found that striatal VMAT2 density is higher in WKY adolescents than in SHR (Cho et al., 2014). Our studies do not show such differences, though a difference between sexes was observed, females had higher levels than males. We reason it may be related to the differential regulation of dopamine transmission in females compared to males (Andersen and Teicher, 2000). However, there are no reports on the sex difference in VMAT2 expression in SHR mice to the best of our knowledge.

The hippocampus is also dysfunctional in ADHD and, conversely, is the brain region most targeted by the positive impact of exercise (Praag et al., 1999; Faraone, 2018). An important molecular mediator of beneficial responses to exercise in the brain is the production of neurotrophins and growth factors, notably BDNF (Park and Poo, 2013) and irisin, an exercise-induced myokine derived from the FNDC5 transmembrane precursor. and acts as a link between skeletal muscle and other tissues (Boström et al., 2012). Interestingly, BDNF and irisin appear to be dysregulated in neuropsychiatric conditions such as depression (REFS) and may comprise links between exercise and brain health (Abdulghani et al., 2023; de Freitas et al., 2020; Islam et al., 2021; Lourenco et al., 2019). In ADHD, irisin facilitates autophagy in the brain possibly acting via glial activation (Abdulghani et al., 2023). Both ADHD subjects and SHR have lower levels of hippocampal BDNF (Fumagalli et al., 2010, 2003; Liu et al., 2015) and physical exercise promoted benefits in the disorder through the upregulation of BDNF (Boström et al., 2012; Seiffer et al., 2022). Our results showed that swimming during pregnancy increased BDNF levels in the hippocampus in male SHR offspring. Our results showed no effects of swimming during pregnancy on irisin in SHR. However, differences between males and females were observed. Females had lower levels of FNDC5/irisin when compared to male rats, which is an intriguing observation that deserves further investigation. To the best our knowledge, there are no studies on the effects of sex on irisin levels in rats and SHR.

In conclusion, the improvement in pain perception and the increase in BDNF levels in the hippocampus in the offspring of SHR rats suggest that swimming during pregnancy has a positive impact that can be leveraged to alleviate symptoms and stimulate neuroplasticity in ADHD.

## Acknowledgements

AT was funded by a fellowship from Coordenação de Aperfeiçoamento de Pessoal de Nível Superior (CAPES) during all the time of her PhD. PP was supported by Fundação Carlos Chagas Filho de Amparo à Pesquisa do Estado Rio de Janeiro (FAPERJ 26/201.430/2021, SEI-260003/004673/2021) and Conselho Nacional de Desenvolvimento Científico e Tecnológico (CNPq 308050/2022-3). MVL was supported by Serrapilheira Institute (R-2012-37967), Alzheimer’s Assocation (Blas Frangione Early Career Achievment Award), FAPERJ (210.316/2022 and 200.248/2023) and CNPq (311487/2019-0 and 309278/2022-8).

## Author contributions

PP and AT designed the study. AT, DM, and PS carried out the behavioral experiments. STF and MVL supervised biochemistry experiments. AT and ASF performed the western blotting and RT-PCR. AT wrote the first draft of the manuscript. STF, ML and PP wrote the final manuscript. All authors contributed to and have approved the final manuscript.

