## Supplemental Tables for "Effects of sex and gestational exercise on pain perception, BDNF and irisin levels in an animal model of ADHD"

**Supplementary Table 1.** Antibodies used in the study

| Antibody | Brand | Reference | Source | Dilution | MW (kDa) |
|---|---|---|---|---|---|
| Antibodies primary | | | | | |
| **TH total** | Milllipore | MAB139 | Mouse | 1:1000 | 55-60 |
| **DAT** | Santa Cruz | SC32258 | Rat | 1:1000 | 55 |
| **D2DR** | Santa Cruz | SC5303 | Mouse | 1:1000 | 50 |
| **FNDC5/Irisin** | Abcam | ab174833 | Rabbit | 1:1000 | 25 |
| **β-actin** | Abcam | ab40390 | Mouse | 1 :2000 | 42 |
| Antibodies secundary | | | | | |
| **Anti-IgG** | Santa Cruz | | Rat | 1:5000 | |
| **Anti-IgG** | Santa Cruz | | Mouse | 1:10000 | |
| **Anti-IgG** | Abcam | | Rabbit | 1 :20000 | |

Table 1. **Supplementary Table 1.** List of antibodies used in quantitative Western blot assays in our study. Abbreviations: TH, tyrosine hydroxylase ; DAT, dopamine transporters;

**Supplementary Table 2.** Primer sequences used in qPCR studies.

| Target | Forward Primer | Reverse Primer |
|---|---|---|
| *TH (r)* | 5’GAGGCTGTCACGTCCCCAA3’ | 5’GGAGGAGGGTTTTGTACCCC3’ |
| *DAT (r)* | 5’CCCCTGCTTCCTCCTGTATG3’ | 5’CACCTCCCCTCTGTCCACTA3’ |
| *VMAT2 (r)* | 5’GACCCTCTAACGTCGCCAAA3’ | 5’CAGCAATGGATGGTGGGACT3’ |
| *D2DR (r)* | 5’GTGCCCTTCATCGTCACTCT3’ | 5’GGTGGGTACAGTTGCCCTTG3’ |
| *ACTB (r)* | 5’TACTGCCCTGGCTCCTAGC3’ | 5’TCAGGAGGAGCAATGATCTTGAT3’ |

Table 2. **Supplementary Table 2.** Primer sequences used in qPCR studies. List of oligonucleotides used in quantitative PCR assays in our study. Abbreviations: TH, tyrosine hydroxylase ; DAT, dopamine transporters ; VMAT2, vesicular monoamine transporter ; Actb, βactin; r, rat.
